# Decoding across sensory modalities reveals common supramodal signatures of conscious perception

**DOI:** 10.1101/115535

**Authors:** Gaëtan Sanchez, Thomas Hartmann, Marco Fuscà, Gianpaolo Demarchi, Nathan Weisz

## Abstract

An increasing number of studies highlight common brain regions and processes in mediating conscious sensory experience. While most studies have been performed in the visual modality, it is implicitly assumed that similar processes are involved in other sensory modalities. However, the existence of supramodal neural processes related to conscious perception has not been convincingly shown so far. Here, we aim to directly address this issue by investigating whether neural correlates of conscious perception in one modality can predict conscious perception in a different modality. In two separate experiments, we presented participants with successive blocks of near-threshold tasks involving tactile, visual or auditory stimuli during the same magnetoencephalography (MEG) acquisition. Using decoding analysis in the post-stimulus period between sensory modalities, our first experiment uncovered supramodal spatio-temporal neural activity patterns predicting conscious perception of the feeble stimulation. Strikingly, these supramodal patterns included activity in primary sensory regions not directly relevant to the task (e.g. neural activity in visual cortex predicting conscious perception of auditory near-threshold stimulation). We carefully replicate our results in a control experiment that furthermore show that the relevant patterns are independent of the type of report (i.e. whether conscious perception was reported by pressing or withholding a button-press). Using standard paradigms for probing neural correlates of conscious perception, our findings reveal a common signature of conscious access across sensory modalities and illustrate the temporally late and widespread broadcasting of neural representations, even into task-unrelated primary sensory processing regions.

## Introduction

While the brain can process an enormous amount of sensory information in parallel, only some information can be consciously accessed, playing an important role in the way we perceive and act in our surrounding environment. An outstanding goal in cognitive neuroscience is thus to understand the relationship between neurophysiological processes and conscious experiences. However, despite tremendous research efforts, the precise brain dynamics that enable certain sensory information to be consciously accessed remain unresolved. Nevertheless, progress has been made in research focusing on isolating neural correlates of conscious perception (1), in particular suggesting that conscious perception - at least if operationalized as reportability (2) - of external stimuli crucially depends on the engagement of a widely distributed brain network (3). To study neural processes underlying conscious perception, neuroscientists often expose participants to near-threshold (NT) stimuli that are matched to their individual perceptual thresholds (4). In NT experiments, there is a trial-to-trial variability in which around 50% of the stimuli at NT-intensity are consciously perceived. Because of the fixed intensity, the physical differences between stimuli within the same modality can be excluded as a determining factor leading to reportable sensation (5). Despite numerous methods used to investigate conscious perception of external events, most studies target a single sensory modality. However, any specific neural pattern identified as a correlate of consciousness needs evidence that it generalizes to some extent, e.g. across sensory modalities. We argue that this has not been convincingly shown so far.

In the visual domain, it has been shown that reportable conscious experience is present when primary visual cortical activity extends towards hierarchically downstream brain areas (6), requiring the activation of frontoparietal regions in order to become fully reportable (7). Nevertheless, a recent MEG study using a visual masking task revealed early activity in primary visual cortices as the best predictor for conscious perception (8). Other studies have shown that neural correlates of auditory consciousness relate to the activation of fronto-temporal rather than fronto-parietal networks (9, 10). Additionally, recurrent processing between primary, secondary somatosensory and premotor cortices have been suggested as potential neural signatures of tactile conscious perception (11, 12). Indeed, recurrent processing between higher and lower order cortical regions within a specific sensory system is theorized to be a marker of conscious processing (6, 13, 14). Moreover, alternative theories such as the global workspace framework (15) extended by Dehaene et al. (16) postulates that the frontoparietal engagement aids in ‘broadcasting’ relevant information throughout the brain, making it available to various cognitive modules. In various electrophysiological experiments, it has been shown that this process is relatively late (∼300 ms), and could be related to increased evoked brain activity after stimulus onset such as the so-called P300 signal (17–19). Such late brain activities seem to correlate with perceptual consciousness and could reflect the global broadcasting of an integrated stimulus making it conscious. Taken together, theories and experimental findings argue in favor of various ‘signatures’ of consciousness from recurrent activity within sensory regions to a global broadcasting of information with engagement of fronto-parietal areas. Even though usually implicitly assumed, it is so far unclear whether similar spatio-temporal neural activity patterns are linked to conscious access across different sensory modalities.

In the current study, we investigated conscious perception in different sensory systems using multivariate analysis on MEG data. Our working assumption is that brain activity related to conscious access has to be independent from the sensory modality: i.e. supramodal consciousness-related neural processes need to exhibit spatio-temporal generalization. Such a hypothesis is most ideally tested applying decoding methods to electrophysiological signals recorded while probing conscious access in different sensory modalities. The application of multivariate pattern analysis (MVPA) to EEG/MEG measurements offers increased sensitivity in detecting experimental effects distributed over space and time (20–23). MVPA is often used in combination with a searchlight method (24, 25), which involves sliding a small spatial window over the data to reveal areas containing decodable information. The combination of both methods provides spatio-temporal detection of optimal decodability, determining where, when and for how long a specific pattern is present in brain activity. Such multivariate decoding analyses have been proposed as an alternative in consciousness research, complementing other conventional univariate approaches in order to identify neural activity predictive of conscious experience at the single trial level (26).

Here, we acquired MEG data while each participant performed three different standard NT tasks on three sensory modalities with the aim of characterizing supramodal brain mechanisms of conscious perception. In the first experiment we show how neural patterns related to perceptual consciousness can be generalized over space and time within and –most importantly-between different sensory systems by using classification analysis on source-level reconstructed brain activity. In an additional control experiment, we replicate the main findings and exclude the possibility that our observed patterns are due to response preparation / selection.

## Materials and Methods

### Participants

Twenty-five healthy volunteers took part in the initial experiment conducted in Trento and twenty-one healthy volunteers took part in the control experiment performed in Salzburg. All participants presented normal or corrected-to-normal vision and no neurological or psychiatric disorders. Three participants for the initial experiment and one participant for the control experiment were excluded from the analysis due to excessive artifacts in the MEG data leading to an insufficient number of trials per condition after artifact rejection (less than 30 trials for at least one condition). Additionally, within each experiment six participants were discarded from the analysis because false alarms rate exceeded 30% and/or near-threshold detection rate was over 85% or below 15% for at least one sensory modality (due to threshold identification failure and difficulty to use response button mapping during the control experiment, also leaving less than 30 trials for at least one relevant condition in one sensory modality: detected or undetected). The remaining 16 participants (11 females, mean age: 28.8 years; SD: 3.4 years) for the initial experiment and 14 participants (9 females, mean age: 26.4 years; SD: 6.4 years) for the control experiment, reported normal tactile and auditory perception. The ethics committee of the University of Trento and University of Salzburg respectively, approved the experimental protocols that were used with the written informed consent of each participant.

### Stimuli

To ensure that the participant did not hear any auditory cues caused by the piezo-electric stimulator during tactile stimulation, binaural white noise was presented during the entire experiment (training blocks included). Auditory stimuli were presented binaurally using MEG-compatible tubal in-ear headphones (SOUNDPixx, VPixx technologies, Canada). Short bursts of white noise with a length of 50 ms were generated with Matlab and multiplied with a Hanning window to obtain a soft on- and offset. Participants had to detect short white noise bursts presented near hearing threshold (27). The intensity of such transient target auditory stimuli was determined prior to the experiment in order to emerge from the background constant white noise stimulation. Visual stimuli were Gabor ellipsoid (tilted 45°) back-projected on a translucent screen by a Propixx DLP projector (VPixx technologies, Canada) at a refresh rate of 180 frames per second. The stimuli were presented 50 ms in the center of the screen at a viewing distance of 110 cm. Tactile stimuli were delivered with a 50 ms stimulation to the tip of the left index finger, using one finger module of a piezo-electric stimulator (Quaerosys, Schotten, Germany) with 2 × 4 rods, which can be raised to a maximum of 1 mm. The module was attached to the finger with tape and the participant’s left hand was cushioned to prevent any unintended pressure on the module (28). For the control experiment (conducted in another laboratory; i.e. Salzburg), visual, auditory and tactile stimulation setups were identical but we used a different MEG/MRI vibrotactile stimulator system (CM3, Cortical Metrics).

### Task and design

The participants performed three blocks of a NT perception task. Each block included three separate runs (100 trials each) for each sensory modality, tactile (T), auditory (A) and visual (V). A short break (∼1 min) separated each run and longer breaks (∼4 min) were provided to the participants after each block. Inside a block, runs alternated in the same order within subject and were pseudo-randomized across subjects (i.e. subject 1 = TVA-TVA-TVA; subject 2 = VAT-VAT-VAT; …). Participants were asked to fixate on a central white dot in a grey central circle at the center of the screen throughout the whole experiment to minimize eye movements.

A short training run with 20 trials was conducted to ensure that participants had understood the task. Then, in three different training sessions prior to the main experiment, participants’ individual perceptual thresholds (tactile, auditory and visual) were determined in the shielded room. For the initial experiment, a 1-up/1-down staircase procedure with two randomly interleaved staircases (one up- and one downward) was used with fixed step sizes. For the control experiment we used a Bayesian active sampling protocol to estimate psychometric slope and threshold for each participant (60).

The main experiment consisted of a detection task (Figure 1A). At the beginning of each run, participants were told that on each trial a weak stimulus (tactile, auditory or visual depending on the run) could be presented at random time intervals. 500 ms after the target stimulus onset, participants were prompted to indicate whether they had felt the stimulus with an on-screen question mark (maximal response time: 2 s). Responses were given using MEG-compatible response boxes with the right index finger and the middle finger (response button mapping was counterbalanced among participants). Trials were then classified into hits (detected) and misses (undetected stimulus) according to the participants’ answers. Trials with no response were rejected. Catch (above perceptual threshold stimulation intensity) and Sham (absent stimulation) trials were used to control false alarms and correct rejection rates across the experiment. Overall, there were 9 runs with 100 trials each (in total 300 trials for each sensory modality). Each trial started with a variable interval (1.3–1.8 s, randomly-distributed) followed by an experimental near-threshold stimulus (80 per run), a sham stimulus (10 per run) or a catch stimulus (10 per run) of 50 ms each. Each run lasted for approximately 5 min. The whole experiment lasted for ∼1h.

**Figure 1.**
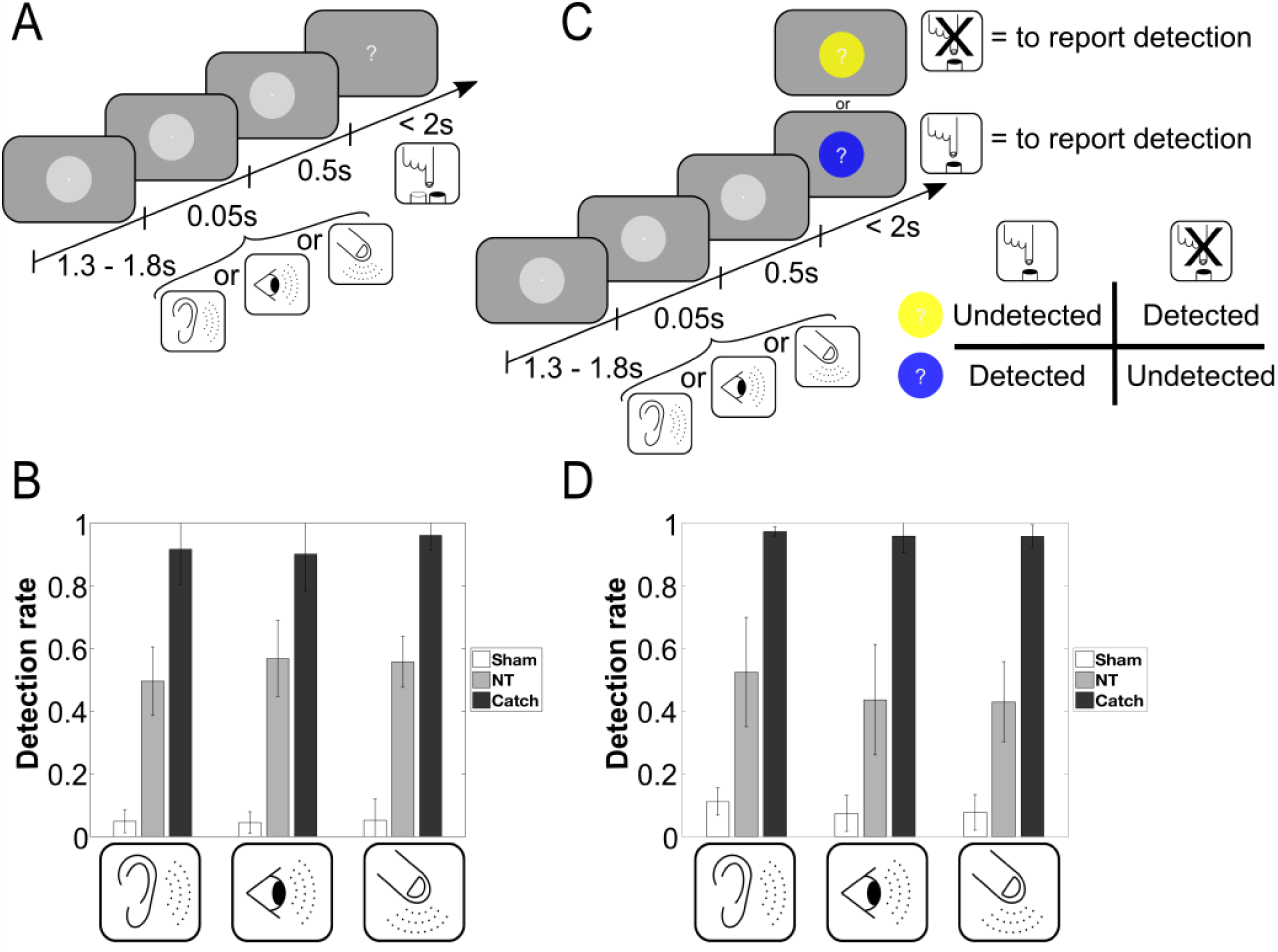
Experimental designs and behavioral results. (**A-B**) Initial experiment; (**C-D**) Control experiment; (**A**) After a variable inter-trial interval between 1.3-1.8 s during which participants fixated on a central white dot, a tactile/auditory/visual stimulus (depending on the run) was presented for 50 ms at individual perceptual intensity. After 500 ms, stimulus presentation was followed by an on-screen question mark, and participants indicated their perception by pressing one of two buttons (i.e. stimulation was ‘present’ or ‘absent’) with their right hand. (**B** & **D**) The group average detection rates for NT stimulation were around 50% across the different sensory modalities. Sham trials in white (no stimulation) and Catch trials in dark (high intensity stimulation) were significantly different from the NT condition in grey within the same sensory modality for both experiments. Error bars depict the standard deviation. (**C**) Identical timing parameters were used in the control experiment; however, a specific response screen design was used to control for motor response mapping. Each trial the participants must use a different response mapping related to circle’s color surrounding the question mark during response screen. Two colors (blue or yellow) were used and presented randomly during the control experiment. One color was associated to the following response mapping rule: “press the button only if there is a stimulation” (for near-threshold condition: “detected”) and the other color was associated to the opposite response mapping: “press a button only if there is no stimulation” (for near-threshold condition: “undetected”). The association between one response mapping and a specific color (blue or yellow) was fixed for a single participant but was predefined randomly across different participant.

Identical timing parameters were used in the control experiment. However, a specific response screen design was used to control for motor response mapping. For each trial the participants must use a different response mapping related to circle’s color surrounding the question mark during response screen. Two colors (blue or yellow) were used and presented randomly after each trial during the control experiment. One color was associated to the following response mapping rule: “press the button only if there is a stimulation” (for near-threshold condition: “detected”) and the other color was associated to the opposite response mapping: “press a button only if there is no stimulation” (for near-threshold condition: “undetected”). The association between one response mapping and a specific color (blue or yellow) was fixed for a single participant but was predefined randomly across different participant. Importantly, by delaying the response-mapping to after the stimulus presentation in a -for the individual-unpredictable manner, neural patterns during relevant periods putatively cannot be confounded by response selection / preparation. Both experiments were programmed in Matlab using the open source Psychophysics Toolbox (61).

### MEG data acquisition and preprocessing

MEG was recorded at a sampling rate of 1kHz using a 306-channel (204 first order planar gradiometers, 102 magnetometers) VectorView MEG system for the first experiment in Trento, and Triux MEG system for the control experiment in Salzburg (Elekta-Neuromag Ltd., Helsinki, Finland) in a magnetically shielded room (AK3B, Vakuumschmelze, Hanau, Germany). Before the experiments, individual head shapes were acquired for each participant including fiducials (nasion, pre-auricular points) and around 300 digitized points on the scalp with a Polhemus Fastrak digitizer (Polhemus, Vermont, USA). Head positions of the individual relative to the MEG sensors were continuously controlled within a run using five coils. Head movements did not exceed 1 cm within and between blocks.

Data were analyzed using the Fieldtrip toolbox (62) and the CoSMoMVPA toolbox (63) in combination with MATLAB 8.5 (MathWorks Natick, MA). First, a high-pass filter at 0.1 Hz (FIR filter with transition bandwidth 0.1Hz) was applied to the continuous data. Then the data were segmented from 1000 ms before to 1000 ms after target stimulation onset and down-sampled to 512 Hz. Trials containing physiological or acquisition artifacts were rejected. A semi-automatic artifact detection routine identified statistical outliers of trials and channels in the datasets using a set of different summary statistics (variance, maximum absolute amplitude, maximum z-value). These trials and channels were removed from each dataset. Finally, the data were visually inspected and any remaining trials and channels with artifacts were removed manually. Across subjects, an average of 5 channels (± 2 SD) were rejected. Bad channels were excluded from the whole data set. A detailed report of remaining number of trials per condition for each participant can be found in supplementary material (see SI Appendix Table S1). Finally, in all further analyses and within each sensory modality for each subject, an equal number of detected and undetected trials was randomly selected to prevent any bias across conditions (64).

### Source analyses

Neural activity evoked by stimulus onset was investigated by computing event-related fields (ERF). For all source-level analyses, the preprocessed data was 30Hz lowpass-filtered and projected to source-level using an LCMV beamformer analysis (65). For each participant, realistically shaped, single-shell headmodels (66) were computed by co-registering the participants’ headshapes either with their structural MRI or – when no individual MRI was available (3 participants and 2 participants, for the initial experiment and the control experiment respectively) – with a standard brain from the Montreal Neurological Institute (MNI, Montreal, Canada), warped to the individual headshape. A grid with 1.5 cm resolution based on an MNI template brain was morphed into the brain volume of each participant. A common spatial filter (for each grid point and each participant) was computed using the leadfields and the common covariance matrix, taking into account the data from both conditions (detected and undetected; or catch and sham) for each sensory modality separately. The covariance window for the beamformer filter calculation was based on 200 ms pre- to 500 ms post-stimulus. Using this common filter, the spatial power distribution was then estimated for each trial separately. The resulting data were averaged relative to the stimulus onset in all conditions (detected, undetected, catch and sham) for each sensory modality. Only for visualization purposes a baseline correction was applied to the averaged source-level data by subtracting a time-window from 200 ms pre-stimulus to stimulus onset. Based on a significant difference between event-related fields of the two conditions over time for each sensory modality, the source localization was performed restricted to specific time-windows of interest. All source images were interpolated from the original resolution onto an inflated surface of an MNI template brain available within the Caret software package (67). The respective MNI coordinates and labels of localized brain regions were identified with an anatomical brain atlas (AAL atlas; (68)) and a network parcellation atlas (29).

### Multivariate Pattern Analysis (MVPA) decoding

MVPA decoding was performed for the period 0 to 500 ms after stimulus onset based on normalized (z-scored) single trial source data downsampled to 100Hz (i.e. time steps of 10 ms). We used multivariate pattern analysis as implemented in CoSMoMVPA (63) in order to identify when and what kind of common network between sensory modality is activated during the near-threshold detection task. We defined two classes for the decoding related to the task behavioral outcome (detected and undetected). For decoding within the same sensory modality, single trial source data were randomly assigned to one of two chunks (half of the original data).

For decoding of all sensory modalities together, single trial source data were pseudo-randomly assigned to one of the two chunks with half of the original data for each sensory modality in each chunk. Data were classified using a 2-fold cross-validation procedure, where a Bayes-Naive classifier predicted trial conditions in one chunk after training on data from the other chunk. For decoding between different sensory modality, single trial source data of one modality were assigned to one testing chunk and the trials from other modalities were assigned to the training chunk. The number of target categories (e.g. detected / undetected) was balanced in each training partition and for each sensory modality. Training and testing partitions always contained different sets of data.

First, the temporal generalization method was used to explore the ability of each classifier across different time points in the training set to generalize to every time point in the testing set (21). In this analysis we used local neighborhoods features in time space (time radius of 10ms: for each time step we included as additional features the previous and next time sample data point). We generated temporal generalization matrices of task decoding accuracy (detected/undetected), mapping the time at which the classifier was trained against the time it was tested. Generalization of decoding accuracy over time was calculated for all trials and systematically depended on a specific between or within sensory modality decoding. The reported average accuracy of the classifier for each time point corresponds to the group average of individual effect-size: the ability of classifiers to discriminate ‘detected’ from ‘undetected’ trials. We summarized time generalization by keeping only significant accuracy for each sensory modalities decoding. Significant classifiers’ accuracies were normalized between 0 and 1:

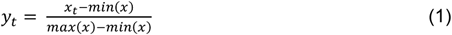

Where *x* is a variable of all significant decoding accuracies and *x*_*t*_ is a given significant accuracy at time *t*. Normalized accuracies (*y*_*t*_) were then averaged across significant testing time and decoding conditions. The number of significant classifier generalization across testing time points and the relevant averaged normalized accuracies were reported along training time dimension (see Figure 3B and 5B). For all significant time points previously identified we performed a ‘searchlight’ analysis across brain sources and time neighborhood structure. In this analysis we used local neighborhoods features in source and time space. We used a time radius of 10ms and a source radius of 3 cm. All significant searchlight accuracy results were averaged over time and only the maximum 10% significant accuracy were reported on brain maps for each sensory modality decoding condition (Figure 4) or for all conditions together (Figure 5C).

**Figure 2.**
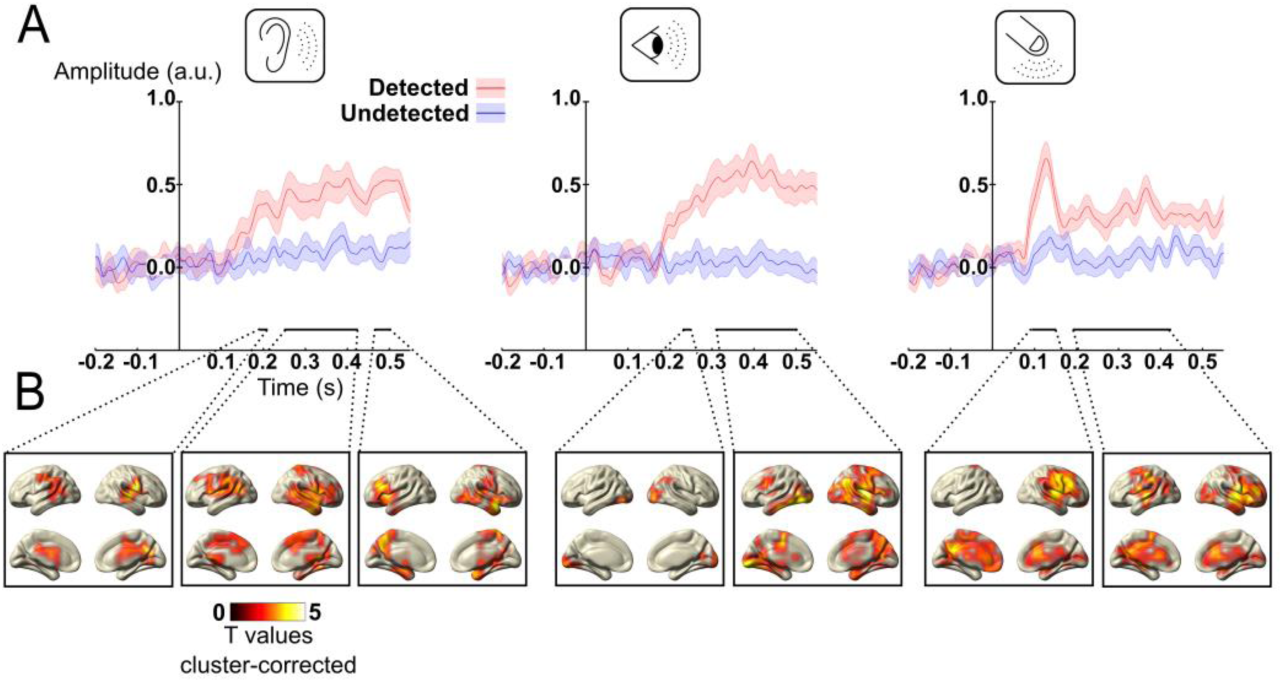
NT trials event-related responses for different sensory modalities: auditory (left panel), tactile (middle panel) and visual (right panel). (**A**) Source-level absolute value (baseline-corrected for visualization purpose) of group event-related average (solid line) and standard error of the mean (shaded area) in the detected (red) and undetected (blue) condition for all brain sources. Significant time windows are marked with bottom solid lines (black line: p_Bonferroni-corrected_ < 0.05) for the contrast detected vs. undetected trials. The relative source localization maps are represented in part B for the averaged time period. (**B**) Source reconstruction of the significant time period marked in part A for the contrast detected vs. undetected trials, masked at p_cluster-corrected_ < 0.05.

**Figure 3.**
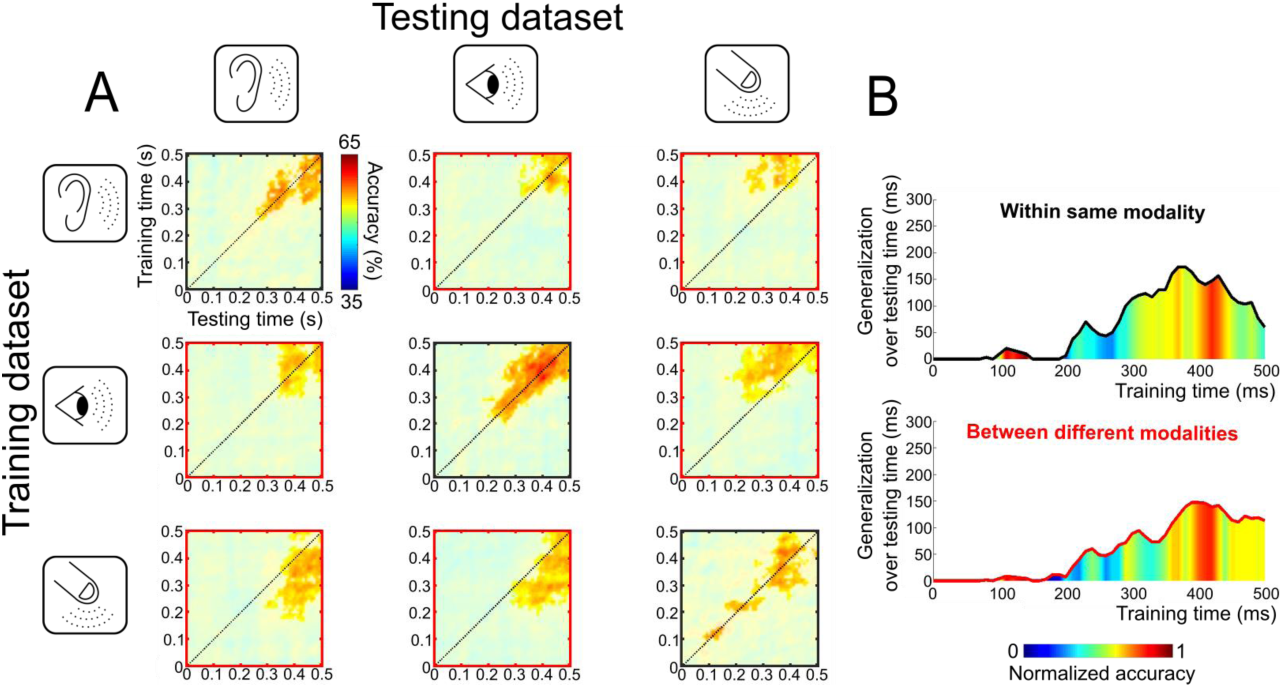
Time-by-time generalization analysis within and between sensory modality (for NT trials). ×3 matrices of decoding results represented over time (from stimulation onset to 500 ms after). **(A)** Each cell presents the result of the searchlight MVPA with time-by-time generalization analysis where classifier accuracy was significantly above chance level (50%) (masked at p_corrected_<0.005). For each temporal generalization matrix, a classifier was trained at a specific time sample (vertical axis: training time) and tested on all time samples (horizontal axis: testing time). The black dotted line corresponds to the diagonal of the temporal generalization matrix, i.e., a classifier trained and tested on the same time sample. This procedure was applied for each combination of sensory modality, i.e. presented on the first row is decoding analysis performed by classifiers trained on the auditory modality and tested on auditory, visual or tactile (1^st^, 2^nd^ and 3^rd^ column respectively) for the two classes: detected and undetected trials. The cells contoured with black line axes (on the diagonal) correspond to within the same sensory modality decoding, whereas the cells contoured with red line axes correspond to between different modalities decoding. **(B)** Summary of average time-generalization and decoding performance over time for all within modality analysis (top panel: average based on the 3 black cells of part A) and between modalities analysis (bottom panel: average based on the 6 red cells of part A). For each specific training time point on the x-axis the average duration of classifier’s ability to significantly generalize on testing time points was computed and reported on the y-axis. Additionally, normalized average significant classifiers accuracies over all testing time for a specific training time point is represented as a color scale gradient.

**Figure 4.**
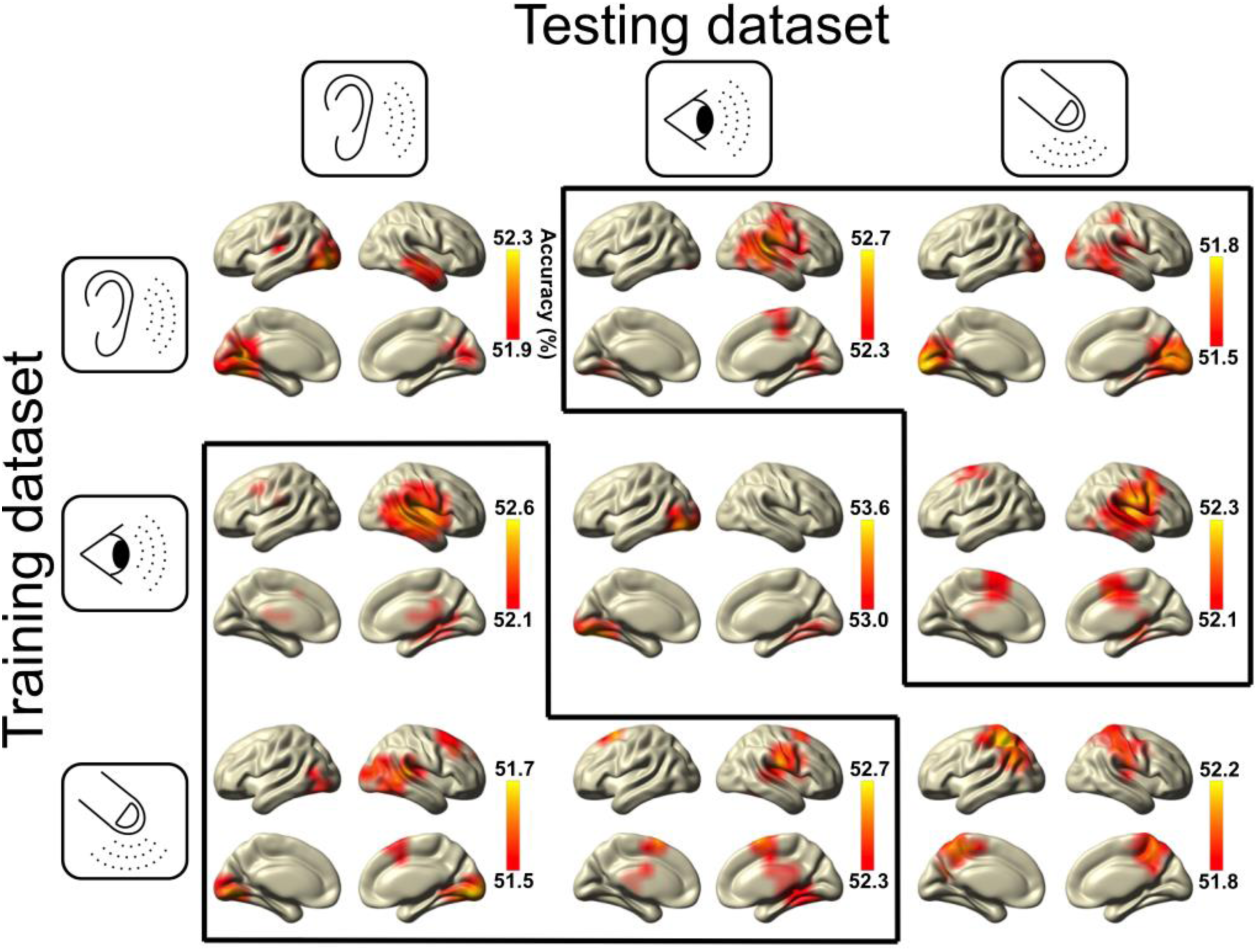
Spatial distribution of significant searchlight MVPA decoding within and between sensory modality. Source brain maps for average decoding accuracy restricted to the related time-generalization significant time-by-time cluster (cf. Figure 3A). Brain maps were thresholded by only showing 10% maximum significant decoding accuracy for each respective time-by-time cluster. Dark solid lines separate all between sensory modality decoding brain maps from the cross-validation within one sensory modality decoding analysis on the diagonal.

**Figure 5.**
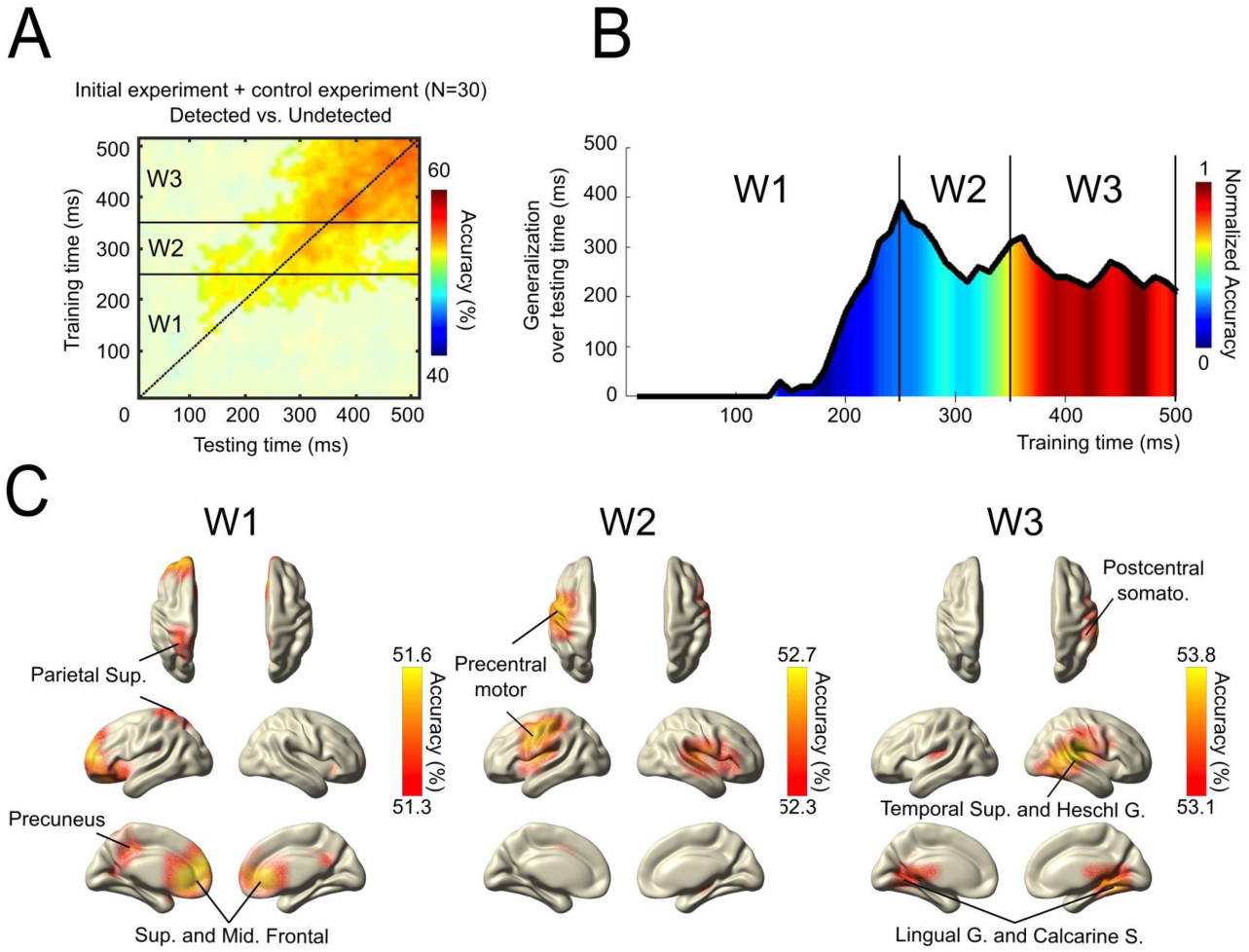
Time-by-time generalization and brain searchlight decoding analysis across all sensory modalities (for NT trials). Compiled results for both initial and control experiments. **(A)** Decoding results represented over time (from stimulation onset to 500 ms after. Result of the searchlight MVPA with time-by-time generalization analysis of “detected” versus “undetected” trials across all sensory modalities. Figure shows the time-clusters where classifier accuracy was significantly above chance level (50%) (masked at p_corrected_<0.005). The black dotted line corresponds to the diagonal of the temporal generalization matrix, i.e., a classifier trained and tested on the same time sample. Horizontal black lines separate time windows (W1, W2 and W3) **(B)** Summary of average time-generalization and decoding performance over time (A). For each specific training time point on the x-axis the average duration of classifier’s ability to significantly generalize on testing time points was computed and reported on the y-axis. Additionally, normalized average significant classifiers accuracies over all testing time for a specific training time point is represented as a color scale gradient. Based on this summary three time windows were depicted to explore spatial distribution of searchlight decoding (W1 : [0 250]ms ; W2 : [250 350]ms ; W3 : [350 500]ms). **(C)** Spatial distribution of significant searchlight MVPA decoding for the significant time clusters depicted in (A) and (B). Brain maps were thresholded by only showing 10% maximum significant (p_corrected_<0.005) decoding accuracy for each respective time-by-time cluster.

Finally, we applied the same type of analysis to all sensory modalities by taking all blocks together with detected and undetected NT trials (equalized within each sensory modality). For the control experiment, we equalized trials based on the 2×2 design with detection report (“detected” or “undetected”) and type of response (“button press = response” or “no response”), so that we get the same number of trials inside each category (i.e. class) for each sensory modality. We performed similar decoding analysis by using different classes definition: either “detected vs. undetected” or “response vs. no response” (SI Appendix, Figure S3B and C).

### Statistical analysis

Detection rates for the experimental trials were statistically compared to those from the catch and sham trials, using a dependent-samples T-Test. Concerning the MEG data, the main statistical contrast was between trials in which participants reported a stimulus detection and trials in which they did not (detected vs. undetected).

The evoked response at the source level was tested at the group level for each of the sensory modalities. To eliminate polarity, statistics were computed on the absolute values of source-level event-related responses. Based on the global average of all grid points, we first identified relevant time periods with maximal difference between conditions (detected vs. undetected) by performing group analysis with sequential dependent T-tests between 0 and 500 ms after stimulus onset using a sliding window of 30 ms with 10ms overlap. P-values were corrected for multiple comparisons using Bonferroni correction. Then, in order to derive the contributing spatial generators of this effect, the conditions ‘detected’ and ‘undetected’ were contrasted for the specific time periods with group statistical analysis using nonparametric cluster-based permutation tests with Monte Carlo randomization across grid points controlling for multiple comparisons (69).

The multivariate searchlight analysis results discriminating between conditions were tested at the group level by comparing the resulting individual accuracy maps against chance level (50%) using a non-parametric approach implemented in CoSMoMVPA (63) adopting 10.000 permutations to generate a null distribution. P-values were set at p<0.005 for cluster level correction to control for multiple comparisons using a threshold-free method for clustering (70), which has been used and validated for MEG/EEG data (38, 71). The time generalization results at the group level were thresholded using a mask with corrected z-score>2.58 (or p_corrected_<0.005) (Figure 3A and 5A). Time points exceeding this threshold were identified and reported for each training data time course to visualize how long time generalization was significant over testing data (Figure 3B and 5B). Significant accuracy brain maps resulting from the searchlight analysis on previously identified time points were reported for each decoding condition. The maximum 10% of averaged accuracies were depicted for each significant decoding cluster on brain maps (Figure 4 and 5).

## Results

### Behavior

We investigated participants’ detection rate for NT, Sham and Catch trials separately for the initial and the control experiment. During the initial experiment participants had to wait for a response screen and press a button on each trial to report their perception (Figure 1A). During the control experiment, however a specific response screen was used to control for motor response mapping. At each trial the participants must use a different response mapping related to circle’s color surrounding the question mark during response screen (see Figure 1C). For the initial experiment and across all participants (N = 16), detection rates for NT experimental trials were: 50% (SD: 11%) for auditory runs, 56% (SD: 12%) for visual runs and 55% (SD: 8%) for tactile runs. The detection rates for the catch trials were 92% (SD: 11%) for auditory runs, 90% (SD: 12%) for visual runs and 96% (SD: 5%) for tactile runs. The mean false alarm rates in sham trials were 4% (SD: 4%) for auditory runs, 4% (SD: 4%) for visual runs and 4% (SD: 7%) for tactile runs (Figure 1B). Detection rates of NT experimental trials in all sensory modality significantly differed from those of catch trials (auditory: T15 = −14.44, p < 0.001; visual: T15 = −9.47, p < 0.001; tactile: T15 = −20.16, p < 0.001) or sham trials (auditory: T15 = 14.66, p < 0.001; visual: T15 = 16.99, p < 0.001; tactile: T15 = 20.66, p < 0.001). Similar results were observed for the control experiment across all participants (N = 14), detection rates for NT experimental trials were: 52% (SD: 17%) for auditory runs, 43% (SD: 17%) for visual runs and 42% (SD: 12%) for tactile runs. The detection rates for the catch trials were 97% (SD: 2%) for auditory runs, 95% (SD: 5%) for visual runs and 95% (SD: 4%) for tactile runs. The mean false alarm rates in sham trials were 11% (SD: 4%) for auditory runs, 7% (SD: 6%) for visual runs and 7% (SD: 6%) for tactile runs (Figure 1B). Detection rates of NT experimental trials in all sensory modality significantly differed from those of catch trials (auditory: T13 = −9.64, p < 0.001; visual: T13 = −10.78, p < 0.001; tactile: T13 = −14.75, p < 0.001) or sham trials (auditory: T13 = 7.85, p < 0.001; visual: T13 = 6.24, p < 0.001; tactile: T13 = 9.75, p < 0.001). Overall the behavioral results are comparable to other studies (27, 28). Individual reaction-times and performances are reported in supplementary materials (see SI Appendix Table S2).

### Event-related neural activity

To compare poststimulus processing for ‘detected’ and ‘undetected’ trials, evoked responses were calculated at the source level for the initial experiment. As a general pattern over all sensory modalities, source-level event-related fields (ERF) averaged across all brain sources show that stimuli reported as detected resulted in pronounced post-stimulus neuronal activity, whereas unreported stimuli did not (Figure 2A). Similar general patterns were observed for the control experiment with identical univariate analysis (see SI Appendix Figure S2). ERFs were significantly different over the averaged time-course with specificity dependent on the sensory modality targeted by the stimulation. Auditory stimulations reported as detected elicit significant differences compared to undetected trials first between 190 and 210 ms, then between 250 and 425ms and finally between 460 and 500 ms after stimulus onset (Figure 2A – left panel). Visual stimulation reported as detected elicits a large increase of ERF amplitude compared to undetected trials from 230-250ms and from 310-500 ms after stimulus onset (Figure 2A – middle panel). Tactile stimulation reported as detected elicits an early increase of ERF amplitude between 95 and 150 ms then a later activation between 190 and 425 ms after stimulus onset (Figure 2A – right panel). Source localization of these specific time periods of interest were performed for each modality (Figure 2B). The auditory condition shows significant early source activity mainly localized to bilateral auditory cortices, superior temporal sulcus and right inferior frontal gyrus, whereas the late significant component was mainly localized to right temporal gyrus, bilateral precentral gyrus, left inferior and middle frontal gyrus. A large activation can be observed for the visual conditions including primary visual areas, fusiform and calcarine sulcus and a large fronto-parietal network activation including bilateral inferior frontal gyrus, inferior parietal sulcus and cingulate cortex. The early contrast of tactile evoked response shows a large difference in the brain activation including primary and secondary somatosensory areas, but also a large involvement of right frontal activity. The late contrast of tactile evoked response presents brain activation including left frontal gyrus, left inferior parietal gyrus, bilateral temporal gyrus and supplementary motor area.

### Decoding and multivariate searchlight analysis across time and brain regions

We investigated the generalization of brain activation over time within and between the different sensory modalities. To this end, we performed a multivariate analysis of reconstructed brain source-level activity from the initial experiment. Time generalization analysis presented as a time-by-time matrix between 0 and 500 ms after stimulus onset shows significant decoding accuracy for each condition (Figure 3A). As can be seen on the black cells located on the diagonal in Figure 3A, cross-validation decoding was performed within the same sensory modality. However, off-diagonal red cells of Figure 3A represent decoding analysis between different sensory modality. Inside each cell, data reported along the diagonal (dashed line) reveal average classifiers accuracy for a specific time point used for the training and testing procedure, whereas off-diagonal data reveal a potential classifier ability to generalize decoding based on different training and testing time points procedure. Indeed, we observed the ability of the same classifier trained on a specific time point to generalize its decoding performance over several time points (see off-diagonal significant decoding inside each cell of Figure 3A). In order to appreciate this result, we computed the average duration of significant decoding on testing time points based on the different training time points (Figure 3B). On average, decoding within the same modality, the classifier generalization starts after 200 ms and we observed significant maximum classification accuracy after 400 ms (see Figure 3B - top panel).

Early differences specific to the tactile modality have been grasped by the classification analysis by showing significant decoding accuracy already after 100 ms without strong time generalization for this sensory modality, where auditory and visual conditions show only significant decoding starting around 250-300 ms after stimulus onset. Such an early dynamic specific to the tactile modality could explain off-diagonal accuracy for all between modalities decoding where the tactile modality was involved (Figure 3A). Interestingly, time generalization analysis concerning between sensory modality decoding (red cells in Figure 3A) revealed significant maximal generalization at around 400 ms (see Figure 3B - bottom panel). In general, the time-generalization analysis revealed time-clusters restricted to late brain activity with maximal decoding accuracy on average after 300 ms for all conditions. The similarity of this time-cluster over all three sensory modalities suggests the generality of such brain activation.

Restricted to the respective significant time clusters (Figure 3A), we investigated the underlying brain sources resulting from the searchlight analysis within and between conditions (Figure 4). The decoding within the same sensory modality revealed higher significant accuracy in relevant sensory cortex for each specific modality condition (see Figure 4; brain plots on diagonal). In addition, auditory modality searchlight decoding revealed also a strong involvement of visual cortices (Figure 4: first row, first column), while somatosensory modality decoding revealed parietal regions involvement such as precuneus (Figure 4: third row, third column). However, decoding searchlight analysis between different sensory modalities revealed higher decoding accuracy in fronto-parietal brain regions in addition to diverse primary sensory regions (see Figure 4; brain plots off diagonal).

### Decoding and multivariate searchlight analysis over all sensory modalities

We further investigated the decoding generalizability of brain activity patterns across all sensory modalities in one analysis by decoding detected versus undetected trials over all blocks together (Figure 5A). Initially, we performed this specific analysis with data from the first experiment and separately with data from the control experiment in order to replicate our findings and control for potential motor response bias (see SI Appendix Figure S3). By delaying the response-mapping to after the stimulus presentation in a random fashion during the control experiment, neural patterns during relevant periods putatively cannot be confounded by response selection / preparation. Importantly, analysis performed on the control experiment used identical data in SI Appendix figure S3 B and C, but only trials assignation (i.e. 2 classes definition) for decoding was different: “detected versus undetected” (SI Appendix, Figure S3B) or “response versus no response” (SI Appendix, Figure S3C). Only decoding of conscious report (i.e. “detected versus undetected”) showed significant time-by- time clusters (SI Appendix, Figure S3 A&B). This result rules out a confounding influence of the motor report and again strongly suggests the existence of a common supramodal pattern related to conscious perception.

We investigated the similarity of time-generalization results by merging data from both experiments (see Figure 5A). We tested for significant temporal dynamics of brain activity patterns across all our data, taking into account that less stable or similar patterns would not survive group statistics. Overall the ability for one classifier to generalize across time seems to increase linearly after a critical time point around 100ms. We show that whereas the early patterns (<250ms) are rather short-lived, temporal generalizability increases showing stability values after ∼350ms (Figure 5B). To follow-up on potential generators underlying these temporal patterns, we depicted the searchlight results from three specific time-windows (W1, W2 and W3) regarding the time-generalization decoding and the distribution of normalized accuracy over time (Figure 5C). W1 from stimulation onset to 250ms depicts the first significant searchlight decoding found in this analysis; W2 from 250ms to 350ms depicts the first generalization period where decoding accuracy is low; finally W3 from 350ms to 500ms depicts the second time-generalization period where higher decoding accuracy were found (Figure 5B). The depiction of the results highlights precuneus, insula, anterior cingulate cortex, frontal and parietal regions mainly involved during the first significant time-window (W1), while the second time-window (W2) main significant cluster is located over left precentral motor cortices. Interestingly the late time-window (W3) shows stronger decoding over primary sensory cortices where accuracy are the highest: lingual and calcarine sulcus, superior temporal and Heschl gyrus and right postcentral gyrus (Figure 5C). The sources depicted by the searchlight analysis, suggest strong overlaps with functional brain networks related to attention and saliency detection (29), especially during the earliest time periods (W1 and W2) (see SI Appendix, Figure S4).

## Discussion

For a neural process to be a strong contender as a neural correlate of consciousness, it should show some generalization e.g. across sensory modalities. This has –despite being implicitly assumed-never been directly tested. To pursue this important issue, we investigated a standard NT experiment targeting three different sensory modalities in order to explore common spatio-temporal brain activity related to conscious perception using multivariate and searchlight analysis. Our findings focusing on the post-stimulus evoked responses are in line with previous studies for each specific sensory modality, showing stronger brain activation when the stimulation was reported as perceived (27, 28, 30). Importantly by exploiting the advantages of decoding, we provide for the first time direct evidence of common electrophysiological correlates of conscious access across sensory modalities.

### ERF time-course differences across sensory modalities

Our first results suggest significant temporal and spatial differences when univariate contrast between ‘detected’ and ‘undetected’ trials were used to investigate sensory-specific evoked responses. At the source level, the global group average activity revealed different significant time periods according to the sensory modality targeted where modulations of evoked responses related to detected trials can be observed (Figure 2A). In the auditory and visual modalities, we found mainly significant differences after 200 ms. In the auditory domain, perception- and attention-modulated sustained responses around 200 ms from sound onset were found in bilateral auditory and frontal regions using MEG (31, 32). Using MEG, a previous study confirmed awareness-related effects from 240 to 500 ms after target presentation during visual presentation (33).

Our results show early differences in the transient responses (for the contrast detected versus undetected) for the somatosensory domain compared to the other sensory modalities, and have been previously identified using EEG at around 100 and 200 ms (34). Moreover, previous MEG studies have shown early brain signal amplitude modulation (<200ms) related to tactile perception in NT tasks (28, 35, 36). Such differences are less pronounced regarding the contrast between catch and sham trials across sensory modality (see SI Appendix Figure S1). Early ERF difference for the tactile NT trials can be due to the experimental setup where auditory and visual targets stimulation emerged from a background stimulation (constant white noise and screen display) whereas tactile stimuli remain isolated transient sensory targets. Despite these differences the time generalization analysis was able to grasp similar brain activity occurring at different time scale across these three sensory modalities.

Source localizations performed with univariate contrasts for each sensory modality suggest differences in network activation with some involvement of similar brain regions in late time windows such as: inferior frontal gyrus, inferior parietal gyrus and supplementary motor area. However, qualitatively similar topographic patterns observed in such analysis cannot easily be interpreted as similar brain processes. The important question is whether these neural activity patterns within a specific sensory modality can be used to decode subjective report of the stimulation within a different sensory context. The multivariate decoding analysis we performed in the next analysis aimed to answer this question.

### Identification of common brain activity across sensory modalities

Multivariate decoding analysis was used to refine spatio-temporal similarity across these different sensory systems. In general, stable characteristics of brain signals have been proposed as a transient stabilization of distributed cortical networks involved in conscious perception (37). Using the precise time resolution of MEG signal and time-generalization analysis, we investigated the stability and time dynamics of brain activity related to conscious perception across sensory systems. The presence of similar brain activity can be revealed between modalities using such a technique, even if significant ERF modulation is distributed over time. As expected, between-modality time-generalization analysis involving tactile runs show off-diagonal significant decoding due to early significant brain activity for the tactile modality (Figure 3A). This result suggests the existence of early but similar brain activity patterns related to conscious perception in the tactile domain compared to auditory and visual modalities.

Generally, decoding results revealed a significant time cluster starting around 300 ms with high classifier accuracy that speaks in favor of a late neural response related to conscious report. Actually, we observed the ability of the same classifier trained on specific time points with a specific sensory modality condition to generalize its decoding performance over several time points with the same or another sensory modality. This result speaks in favor of supramodal brain activity patterns that are consistent and stable over time. In addition, the searchlight analysis across brain regions provides an attempt to depict brain network activation during these significant time-generalization clusters. Note that, as seen also in multiple other studies using decoding (22, 23, 38, 39), the average accuracy can be relatively low and yet remains significant at the group level. Note however that contrary to many other cognitive neuroscientific studies using decoding (39, 40), we do not apply the practice of “subaveraging” trials to create “pseudo”-single trials, which naturally boosts average decoding accuracy (41). Also, the statistical rigor of our approach is underlined by the fact that the reported decoding results are restricted to highly significant effects (P_corrected_<0.005; see Methods section). Critically, we replicated our results -applying the identical very conservative statistical thresholds-within a second control experiment when looking at conscious perception report contrast independently from motor response activity (SI Appendix, Figure S3). Our results conform to those of previous studies in underlying the importance of late activity patterns as crucial markers of conscious access (7, 42) and decision-making processes (10, 43).

Furthermore in this study, we explored the brain regions underlying time dynamics of conscious report by using brain source searchlight decoding. Knowing the limitations of such MEG analysis, especially using low spatial resolution (3cm), we restricted depiction of results to the main 10% maximum decoding accuracy over all searchlight brain regions. Some of the brain regions found in our searchlight analysis, namely deep brain structures such as the insula and anterior cingulate cortex are shared with other functional brain networks such as the salience network (44, 45). Also the superior frontal and parietal cortex have been previously found to be activated by attention-demanding cognitive tasks (46). Hence, we would like to emphasize that one cannot conclude from our study that the observed network identified in figure 5C is exclusively devoted to conscious report. Brain networks identified in this study share common brain regions and dynamics with the attentional and salience networks that remain relevant mechanisms to performing a NT-task. Interestingly this part of the network seems to be more involved during the initial part of the process, prior to motor brain region involvement (Figure 5C and SI Appendix Figure S4).

Indeed, some brain regions involved in motor planning were identified with our analysis, such as precentral gyrus, and could in principle relate to the upcoming button-press to report the subjective perception of the stimulus. We specifically targeted such motor preparation bias within the control experiment, in which the participant was unable to predict a priori how to report a conscious percept (i.e. pressing or withholding a button press) until the response prompt appeared. Importantly, we did not find any significant decoding when trials used for the analysis where sorted under response type (e.g. with or without an actual button press from the participant) compared to subjective report of detection (see SI Appendix, Figure S3 B and C). Such findings could speak in favor of generic motor planning (47) or decision processes related activity in such forced-choice paradigms (48, 49).

### Late involvement of all primary sensory cortices

Some within-modalities decoding results highlighted unspecific primary cortices involvement while decoding was performed on another sensory modality. For instance, during auditory near-threshold stimulation, the main decoding accuracy of neural activity predicting conscious perception was found in auditory cortices but also in visual cortices (see Figure 4: first row, first column). Interestingly, our final analysis revealed and confirmed that primary sensory regions are strongly involved in decoding conscious perception across sensory modalities. Moreover, such brain regions were mainly found during the last time period investigated following the first main involvement of fronto-parietal areas (see Figure 5). These important results suggest that sensory cortices from a specific modality contain sufficient information to allow the decoding perceptual conscious access in another different sensory modality. These results suggest a late active role of primary cortices over three different sensory systems (Figure 5). One study reported efficient decoding of visual object categories in early somatosensory cortex using fMRI and multivariate pattern analysis (50). Another fMRI experiment suggested that sensory cortices appear to be modulated via a common supramodal frontoparietal network, attesting to the generality of attentional mechanism toward expected auditory, tactile and visual information (51). However, in our study we demonstrate how local brain activity from different sensory regions reveal a specific dynamic allowing generalization over time to decode the behavioral outcome of a subjective perception in another sensory modality. These results speak in favor of intimate cross-modal interactions between modalities in perception (52).

Finally, our results suggest that primary sensory regions remain important at late latency after stimulus onset for resolving stimulus perception over different sensory modalities. We propose that this network could enhance the processing of behaviorally relevant signals, here the sensory targets. Although the integration of classically unimodal primary sensory cortices into a processing hierarchy of sensory information is well established (53), some studies suggest multisensory roles of primary cortical areas (54, 55).

Today it remains unknown how such multisensory responses could be related to an individual’s unisensory conscious percepts in humans. Since sensory modalities are usually interwoven in real life, our findings of a supramodal network that may subserve both conscious access and attentional functions have a higher ecological validity than results from previous studies on conscious perception for single sensory modality.

Actually, our results are in line with an ongoing debate in neuroscience asking to what extent multisensory integration emerges already in primary sensory areas (55, 56). Animal studies provided compelling evidence suggesting that the neocortex is essentially multisensory (57). Here our findings speak in favor of a multisensory interaction in primary and associative cortices. Interestingly a previous an fMRI study by using multivariate decoding revealed distinct mechanisms governing audiovisual integration in primary and associative cortices needed for spatial orienting and interactions in a multisensory world (58).

### Conclusion

We successfully characterized common patterns over time and space suggesting generalization of consciousness-related brain activity across different sensory NT tasks. Our study paves the way for future investigation using techniques with more precise spatial resolution such as functional magnetic resonance imaging to depict in detail the brain network involved. However, to our knowledge this is the first study to report significant spatio-temporal decoding across different sensory modalities near-threshold perception experiment. Indeed, our results speak in favor of the existence of stable and supramodal brain activity patterns, distributed over time and involving seemingly task-unrelated primary sensory cortices. The stability of brain activity patterns over different sensory modalities presented in this study is, to date, the most direct evidence of a common network activation leading to conscious access (2). Moreover, our findings add to recent remarkable demonstrations of applying decoding and time generalization methods to MEG (21–23, 59), and show a promising application of MVPA techniques to source level searchlight analysis with a focus on the temporal dynamics of conscious perception.

## Acknowledgements

This work was supported by the European Research Council (WIN2CON, ERC StG 283404). We thank Julia Frey for her great support during data collection.

## Author contributions

G.S. and N.W. conceived the approach. G.S., G.P. and T.H. implemented the experiment. G.S. and M.F. collected the data. G.S. analyzed the data. G.S. and N.W. wrote the manuscript. All authors approved the current manuscript.

## Resource sharing and data availability

Further information and requests for resources or data should be directed to and will be fulfilled by the corresponding author.

