## Supplementary material for "Decoding across sensory modalities reveals common supramodal signatures of conscious perception": SI Appendix

Corresponding author: Gaëtan Sanchez

**This PDF file includes:**

Figs. S1 to S4
Tables S1 to S2

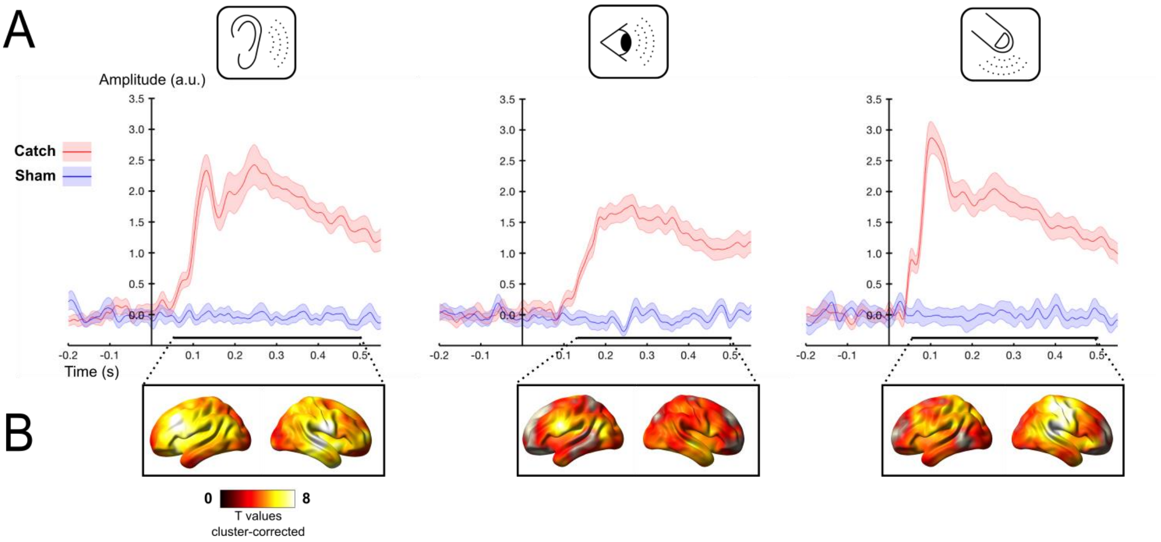


**Fig. S1. Initial experiment. Catch and Sham trials event-related responses for different sensory modalities: auditory (left panel), tactile (middle panel) and visual (right panel).** (A) Group event-related average of all brain sources absolute value average activity (solid line) and standard error of the mean (shaded area) for catch trials (red) and sham trials (blue) condition. Significant time windows are marked with bottom solid lines (black line: $p_{Bonferroni\text{-}corrected} < 0.05$) for the contrast catch vs. sham trials. The relative source localization maps are represented in part B for the average time period. (B) Significant source activity of the average time period marked in part A for the contrast catch vs. sham trials, masked at $p_{cluster\text{-}corrected} < 0.05$.

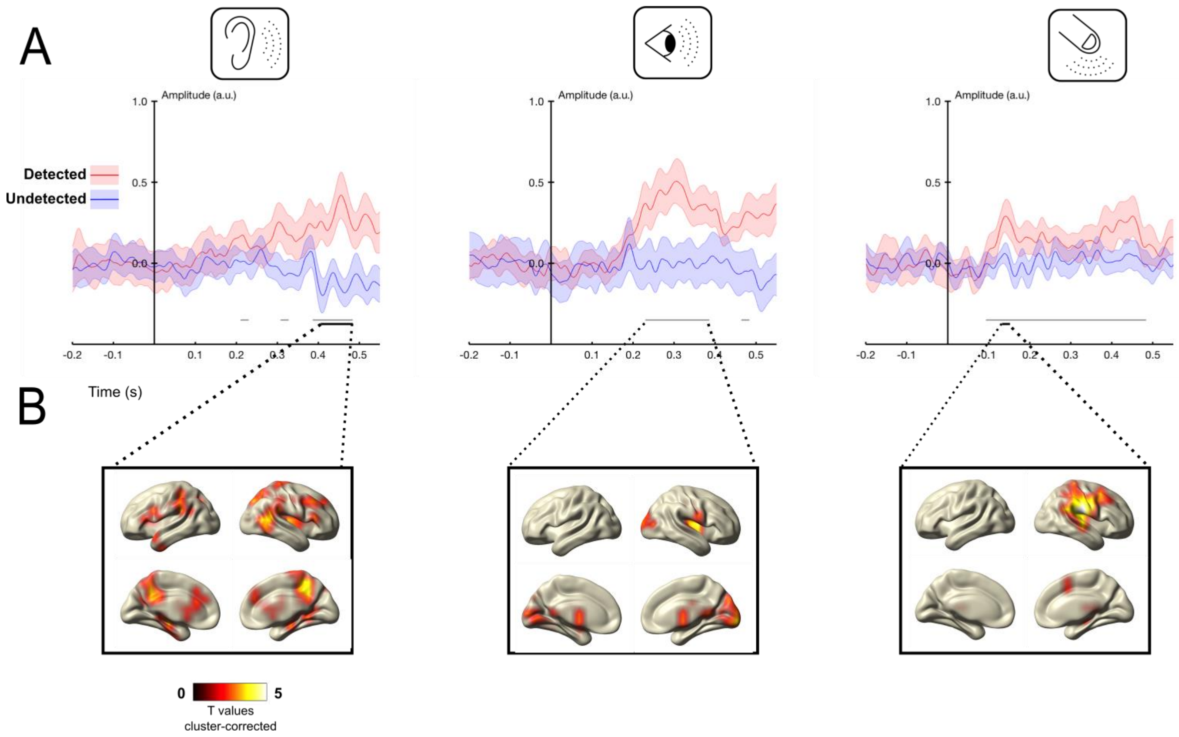


**Fig. S2. Control experiment. NT trials event-related responses for different sensory modalities: auditory (left panel), tactile (middle panel) and visual (right panel).** (A) Source-level absolute value of group event-related average (solid line) and standard error of the mean (shaded area) in the detected (red) and undetected (blue) condition for all brain sources. Significant time windows are marked with bottom solid lines (black line: $p_{bonferroni\text{-}corrected} < 0.05$; grey line: $p_{uncorrected} < 0.05$) for the contrast detected vs. undetected trials. The relative source localization maps are represented in part B for the averaged time period. (B) Source reconstruction of the significant time period marked in part A for the contrast detected vs. undetected trials, masked at $p_{cluster\text{-}corrected} < 0.05$.

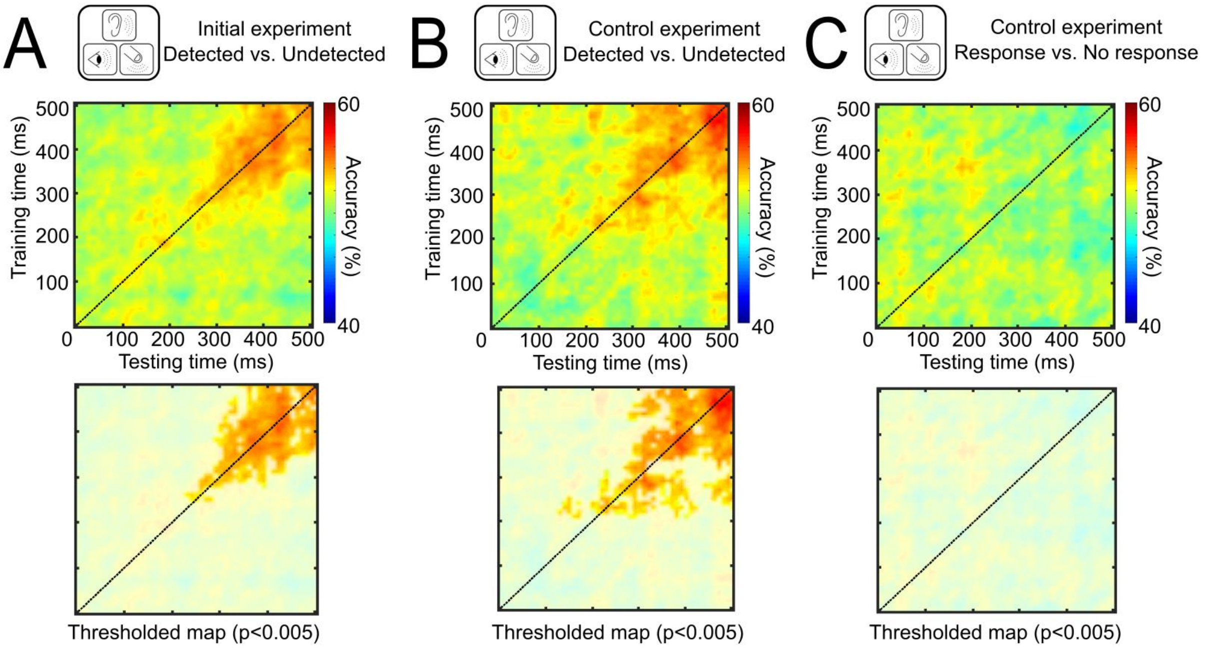


**Figure S3**. **Time-by-time generalization and brain searchlight decoding analysis across all sensory modalities (for NT trials).** Decoding results represented over time (from stimulation onset to 500 ms after). **(A) Initial experiment**, first row shows result of the searchlight MVPA with time-by-time generalization analysis of “detected” versus “undetected” trials across all sensory modalities. Second row shows the time-clusters where classifier accuracy was significantly above chance level (50%) (masked at $p_{corrected}<0.005$). The black dotted line corresponds to the diagonal of the temporal generalization matrix, i.e., a classifier trained and tested on the same time sample. **(B) Control experiment**, similar analysis and plots regarding the contrast “detected” versus “undetected” by using identical number of trials with counterbalancing number of absence or presence of motor response for NT trials. **(C) Control experiment**, similar analysis and plots regarding the contrast “response” versus “no response” by using identical number of trials with counterbalancing number of detected or undetected target stimulation for NT trials.

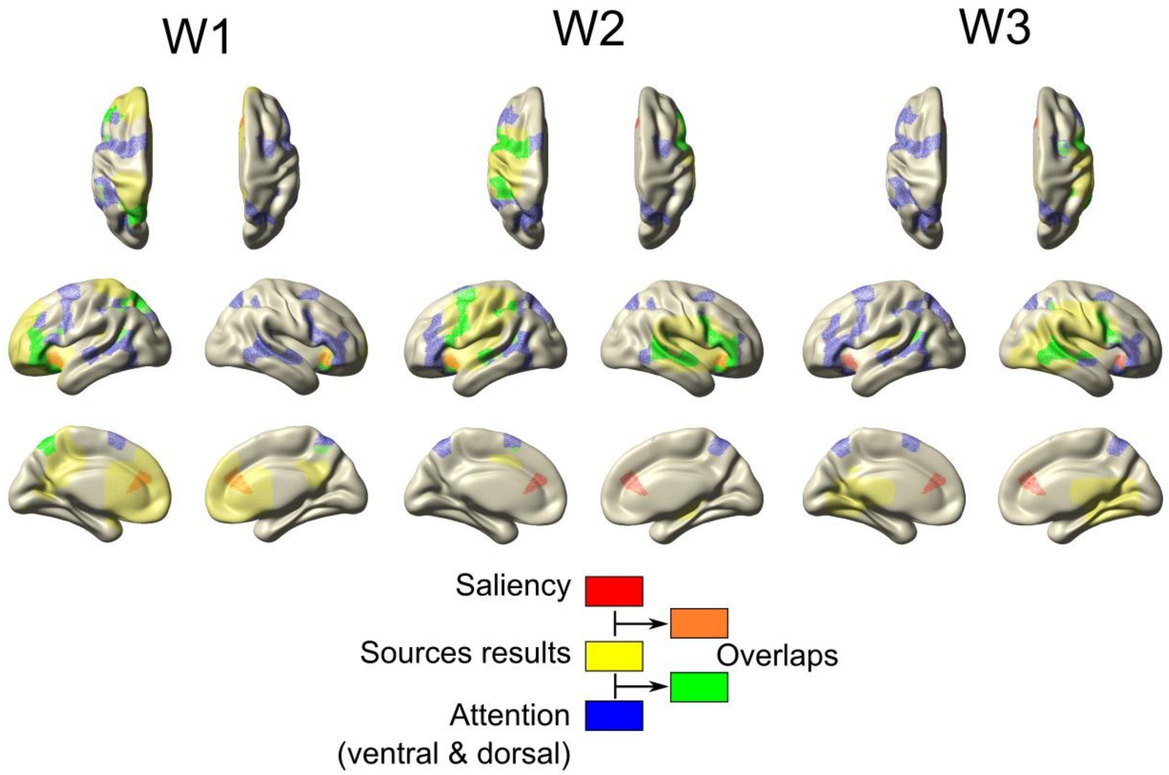


**Figure S4**. **Sources results overlapping representation with attention and saliency network parcels** (1)**.** Compiled results for both initial and control experiments (see figure 5) based on a summary results from three time windows depicted to explore spatial distribution of searchlight decoding (W1 : [0 250]ms ; W2 : [250 350]ms ; W3 : [350 500]ms). Saliency network parcels located over anterior cingulate gyrus and insula regions were found overlapping with maximum searchlight decoding over early time period (W1). Attentional network parcels are maximally overlapping with sources results of the study during W1 and W2. However W3 time period results show less overlap with saliency and attention network parcels due to a spatial distribution of searchlight analysis mainly located in primary sensory areas.

**Table S1. Report of stimulation intensity and equal number of trials analyzed per condition used in decoding analysis for each experiment.** Calibration of near-threshold stimulation intensity is dependent on the participant and the sensory modality targeted: (AUD) auditory stimulation intensity is expressed in dB compared to the background white noise; (VIS) visual stimulation intensity is expressed as normalized arbitrary unit of Gabor patch brightness on the screen; (TAC) tactile stimulation is expressed as arbitrary unit dependent on the tactile stimulator. For each condition we equalized and reported the number of trials for each participant, sensory modality and target conditions. The number reported mean that we used this amount of trials for each condition to conduct our analysis (D/U = detected versus undetected; C/S = catch versus sham; U/D/R/noR = "detected with or without response" versus "undetected with or without response" and "Response for detected or undetected" versus "No response for detected or undetected").

| | | **Stimulation intensity** | | | **Equal number of trials per condition** | | | | | | | | |
|---|---|---|---|---|---|---|---|---|---|---|---|---|---|
| | | **AUD** | **VIS** | **TAC** | **AUD** | | | **VIS** | | | **TAC** | | |
| **Experiment** | **Subject** | | | | **D/U** | **C/S** | **D/U/R/noR** | **D/U** | **C/S** | **D/U/R/noR** | **D/U** | **C/S** | **D/U/R/noR** |
| Initial | 1 | 1,8 | 1,3 | 539 | 109 | 21 | - | 108 | 28 | - | 88 | 29 | - |
| | 2 | 2,3 | 1,3 | 508 | 86 | 26 | - | 84 | 24 | - | 82 | 27 | - |
| | 3 | 1,7 | 1,1 | 549 | 148 | 24 | - | 68 | 26 | - | 45 | 25 | - |
| | 4 | 1,8 | 0,8 | 498 | 74 | 17 | - | 130 | 29 | - | 99 | 27 | - |
| | 5 | 1,8 | 1,9 | 852 | 69 | 25 | - | 123 | 26 | - | 93 | 25 | - |
| | 6 | 1,8 | 1,3 | 593 | 131 | 18 | - | 77 | 25 | - | 65 | 22 | - |
| | 7 | 1,8 | 1,6 | 701 | 137 | 25 | - | 69 | 23 | - | 94 | 24 | - |
| | 8 | 1,8 | 1,6 | 644 | 102 | 27 | - | 125 | 27 | - | 50 | 25 | - |
| | 9 | 1,9 | 1,0 | 683 | 104 | 25 | - | 108 | 25 | - | 70 | 28 | - |
| | 10 | 1,8 | 1,2 | 477 | 78 | 18 | - | 90 | 23 | - | 81 | 25 | - |
| | 11 | 1,7 | 1,2 | 607 | 106 | 29 | - | 84 | 29 | - | 98 | 23 | - |
| | 12 | 1,8 | 5,2 | 525 | 112 | 27 | - | 109 | 27 | - | 83 | 28 | - |
| | 13 | 1,7 | 1,2 | 609 | 108 | 28 | - | 114 | 27 | - | 76 | 25 | - |
| | 14 | 2,2 | 0,9 | 569 | 97 | 29 | - | 129 | 23 | - | 67 | 24 | - |
| | 15 | 1,9 | 1,6 | 675 | 105 | 27 | - | 113 | 23 | - | 113 | 25 | - |
| | 16 | 1,8 | 1,4 | 617 | 125 | 30 | - | 102 | 27 | - | 52 | 27 | - |
| Control | 1 | 5,2 | 6,4 | 8,2 | 89 | 23 | 41 | 38 | 24 | 17 | 88 | 21 | 43 |
| | 2 | 5,9 | 2,2 | 17,1 | 92 | 25 | 42 | 56 | 25 | 19 | 60 | 26 | 27 |
| | 3 | 3,7 | 2,0 | 15,5 | 85 | 25 | 38 | 96 | 20 | 45 | 83 | 24 | 36 |
| | 4 | 0,2 | 1,3 | 10,8 | 81 | 23 | 39 | 51 | 27 | 25 | 98 | 31 | 47 |
| | 5 | 0,4 | 1,1 | 8,5 | 72 | 23 | 32 | 105 | 25 | 52 | 70 | 37 | 33 |
| | 6 | 0,2 | 1,8 | 13,0 | 71 | 18 | 32 | 71 | 22 | 33 | 72 | 20 | 34 |
| | 7 | 0,5 | 1,2 | 11,0 | 96 | 23 | 46 | 102 | 21 | 45 | 91 | 23 | 38 |
| | 8 | 1,9 | 1,2 | 14,1 | 69 | 28 | 33 | 89 | 22 | 39 | 105 | 23 | 49 |
| | 9 | 0,1 | 0,8 | 9,0 | 92 | 23 | 44 | 33 | 20 | 16 | 82 | 22 | 39 |
| | 10 | 1,3 | 2,0 | 13,0 | 96 | 27 | 46 | 86 | 29 | 36 | 65 | 27 | 30 |
| | 11 | 1,9 | 1,3 | 22,6 | 63 | 19 | 27 | 73 | 22 | 35 | 75 | 21 | 36 |
| | 12 | 0,4 | 1,9 | 12,4 | 30 | 17 | 12 | 80 | 19 | 36 | 74 | 17 | 30 |
| | 13 | 1,0 | 0,9 | 5,9 | 58 | 33 | 29 | 65 | 27 | 32 | 108 | 28 | 53 |
| | 14 | 1,9 | 8,0 | 14,2 | 80 | 26 | 30 | 83 | 25 | 31 | 69 | 32 | 30 |

**Table S2. Reaction time and d' values for each participant and conditions.** The control experiment was more challenging on trial-by-trial basis because response mapping was shuffled. Thus false alarm rate and reaction time increased compared to the initial experiment.

| | | d' | | | Reaction time (ms) | | |
|---|---|---|---|---|---|---|---|
| **Experiment** | **Subject** | **AUD** | **VIS** | **TAC** | **AUD** | **VIS** | **TAC** |
| Initial | 1 | 2,0 | 2,2 | 2,3 | 327 | 282 | 304 |
| | 2 | 2,1 | 1,5 | 2,2 | 394 | 442 | 396 |
| | 3 | 2,2 | 1,8 | 2,0 | 342 | 292 | 332 |
| | 4 | 1,4 | 1,4 | 2,0 | 439 | 373 | 539 |
| | 5 | 1,6 | 1,9 | 2,3 | 467 | 399 | 497 |
| | 6 | 2,4 | 2,0 | 0,8 | 286 | 231 | 346 |
| | 7 | 2,0 | 1,5 | 2,1 | 344 | 310 | 453 |
| | 8 | 2,0 | 2,6 | 2,3 | 382 | 322 | 345 |
| | 9 | 1,0 | 2,6 | 2,1 | 402 | 358 | 383 |
| | 10 | 1,1 | 2,3 | 1,9 | 436 | 291 | 433 |
| | 11 | 2,1 | 1,8 | 2,2 | 382 | 357 | 375 |
| | 12 | 1,6 | 2,3 | 2,4 | 447 | 402 | 455 |
| | 13 | 1,8 | 2,2 | 2,1 | 785 | 620 | 835 |
| | 14 | 0,9 | 1,2 | 1,9 | 682 | 703 | 792 |
| | 15 | 2,1 | 2,0 | 1,0 | 536 | 513 | 592 |
| | 16 | 1,6 | 2,2 | 2,1 | 430 | 376 | 394 |
| **Mean** | | 1,7 | 2,0 | 2,0 | 443 | 392 | 467 |
| **STD** | | 0,5 | 0,4 | 0,5 | 130 | 126 | 156 |
| control | 1 | 0,7 | 0,1 | 0,4 | 690 | 718 | 680 |
| | 2 | 1,7 | 2,5 | 1,9 | 572 | 552 | 630 |
| | 3 | 1,8 | 1,6 | 1,5 | 710 | 676 | 722 |
| | 4 | 1,4 | 2,8 | 1,1 | 706 | 604 | 761 |
| | 5 | 1,9 | 1,4 | 1,4 | 681 | 663 | 790 |
| | 6 | 1,6 | 1,9 | 1,3 | 641 | 635 | 750 |
| | 7 | 1,0 | 1,3 | 1,7 | 619 | 670 | 679 |
| | 8 | 1,6 | 0,9 | 1,8 | 704 | 705 | 726 |
| | 9 | 1,0 | 0,1 | 1,4 | 694 | 704 | 676 |
| | 10 | 2,0 | 2,4 | 1,3 | 506 | 533 | 574 |
| | 11 | 0,6 | 1,5 | 1,3 | 732 | 671 | 704 |
| | 12 | 0,1 | 0,5 | 0,7 | 913 | 754 | 851 |
| | 13 | 2,0 | 1,6 | 1,9 | 615 | 731 | 697 |
| | 14 | 1,1 | 1,2 | 1,0 | 723 | 634 | 741 |
| **Mean** | | 1,3 | 1,4 | 1,3 | 679 | 661 | 713 |
| **STD** | | 0,6 | 0,8 | 0,4 | 93 | 64 | 68 |
